# ATFS-1 regulates peroxisome assembly genes and protects both mitochondria and peroxisomes during peroxin perturbations

**DOI:** 10.1101/2022.09.09.507280

**Authors:** Tomer Shpilka, Nandhitha Uma Naresh, YunGuang Du, Jennifer L. Watts, Cole M. Haynes

## Abstract

Peroxisome biogenesis disorders lead to a myriad of clinical manifestations, among which is the dysfunction of the mitochondria. Mitochondria dysfunction is typically sensed by the UPR^mt^, a broad protective transcriptional response governed by the transcription factor ATFS-1. Here, we investigated the role of the UPR^mt^ during peroxisomal stress. We show that mutations or knockdown of peroxins, the genes required for peroxisome assembly, lead to mitochondria dysfunction and the induction of the UPR^mt^ in *C.elegans*. The UPR^mt^ induced the transcription of the mitochondrial outer membrane translocase *mspn-1* (ATAD-1), that in turn alleviated mitochondrial stress most likely by extracting mislocalized proteins. Importantly, ATFS-1 regulated the transcription of peroxins and the peroxisomal transporter *pmp-4*. A *prx-5* loss of function strain induced a retrograde response that resulted in the transcriptional induction of peroxins, peroxisomal transporters, chaperons and proteases. And was dependent on ATFS-1 and on the peroxisome proliferator activator receptor alpha, NHR-49. Knockout of *atfs-1* during peroxisomal stress resulted in severe developmental delays, import defects to peroxisomes and the appearance of large peroxisomal structures. We propose that ATFS-1 regulates the biogenesis of peroxisomes and protects the organism by alleviating the stress of both peroxisomes and mitochondria during peroxisomal stress.

## Introduction

Peroxisomes are ubiquitous organelles, involved in many biochemical processes including fatty acid metabolism, bile acid synthesis and the metabolism of reactive oxygen species^1^. Given their important role in metabolism, peroxisome biogenesis is a highly regulated process, orchestrated by a set of peroxisome assembly factors known as peroxins (PEX/PRX)^2^.

Most of the peroxisomal matrix proteins contain a targeting signal 1 (PTS1) that is recognized by the cytosolic receptor PEX5/PRX-5, which shuttles the proteins to the peroxisome^3–5^. Mutations in *pex5* and in most of the *pex* genes, lead to peroxisome biogenesis disorders that can be mild, moderate or result in the complete loss of peroxisomes. The most severe form, known as the Zellweger Spectrum Disorder (ZSD), lead to a vast array of defects in organelles such as the brain, liver and kidneys as well as difficulties in moving and feeding. There is no cure or treatment to ZSD^6–8^.

Several studies have shown that mutations or deletions in peroxins lead to a reduction in mitochondrial membrane potential and respiration^9,10^. This was attributed to the mislocalization of peroxisomal proteins to the mitochondria and was shown to be alleviated by over expressing the AAA+ ATPase ATAD-1 (MSPN-1 in worms)^11^, which extracts the mislocalized proteins^12,13^.

Reduction in mitochondrial membrane potential and respiration are known inducers of the mitochondrial unfolded response (UPR^mt^)^14,15^, a broad transcriptional response governed by the transcription factor ATFS-1^14,16^. Activation of the UPR^mt^ leads to mitochondria biogenesis^17^, metabolic alterations^18,19^ and the induction of numerous genes that provide protection to the organism^17,19^. Notably, induction of the UPR^mt^ is not always beneficial to the organism and can have detrimental effects. For example, when the UPR^mt^ is induced due to mutations in mitochondrial DNA, the UPR^mt^ leads to the propagation and maintenance of the deleterious mitochondrial DNA^20,21^. Therefore, it is important to understand the contribution of the UPR^mt^ under different forms of stress.

Here we investigated the role of the UPR^mt^ during peroxisomal stress in *C.elegans*. We show that the UPR^mt^ is induced in response to mutations or knockdowns of peroxins and lead to the transcriptional upregulation of *mspn-1*. Strikingly, the UPR^mt^ regulated a set of peroxins required for peroxisomal biogenesis. A combination of *prx-5* loss of function with *atfs-1* knockout resulted in severe developmental delays, reduced import to peroxisomes and the appearance of large peroxisomal structures. Mutations in the peroxin *prx-5* induced a retrograde response that was dependent on ATFS-1 and the peroxisome proliferator activator receptor alpha, NHR-49^22^. Finally, we show that *nhr-49* and *atfs-1* genetically interact. We propose that the ATFS-1 regulates peroxisome assembly and orchestrates the biogenesis of both the peroxisomes and the mitochondria.

## Results

### Peroxin defects or knockdown lead to mitochondria dysfunction and induce the UPR^mt^

To determine whether peroxin defects lead to mitochondrial dysfunction in the model organism *C. elegans*, we utilized a partial loss of function mutant strain of the essential peroxin gene *prx-5* (strain *ku517*), which is required to transport proteins to the peroxisomal matrix^23,24^. This strain harbors a point mutation in the C-terminus of *prx-5*,leading to an early stop codon that reduces the function of the protein, while maintaining worm development, which is otherwise arrested in peroxin deficient strains^25^. To assess mitochondrial function in the *prx-5* mutant strain, we stained the worms with the membrane potential dependent dye Tetramethylrhodamine, Ethyl Ester (TMRE). TMRE staining indicated that the *prx-5(ku517*) strain harbored fewer functional mitochondria than WT worms (Fig. 1*a-b*). And resulted in significant reductions in basal oxygen consumption as well as in the maximal respiratory capacity of the worms (Fig. 1*c-d*). In agreement with these results, RNAi targeted against different peroxins resulted in reduction of TMRE signal (Fig. 1*e*) and in fragmentation of the mitochondrial network in muscle cells (Fig. 1*f*). These phenotypes are consistent with those observed in cells derived from Zellweger patients and mouse models that lack one of the peroxin genes^9–11,26^. Thus, we conclude that *C. elegans* is a suitable model organism to investigate mitochondrial perturbations derived from peroxisomal defects.

**Figure 1:**
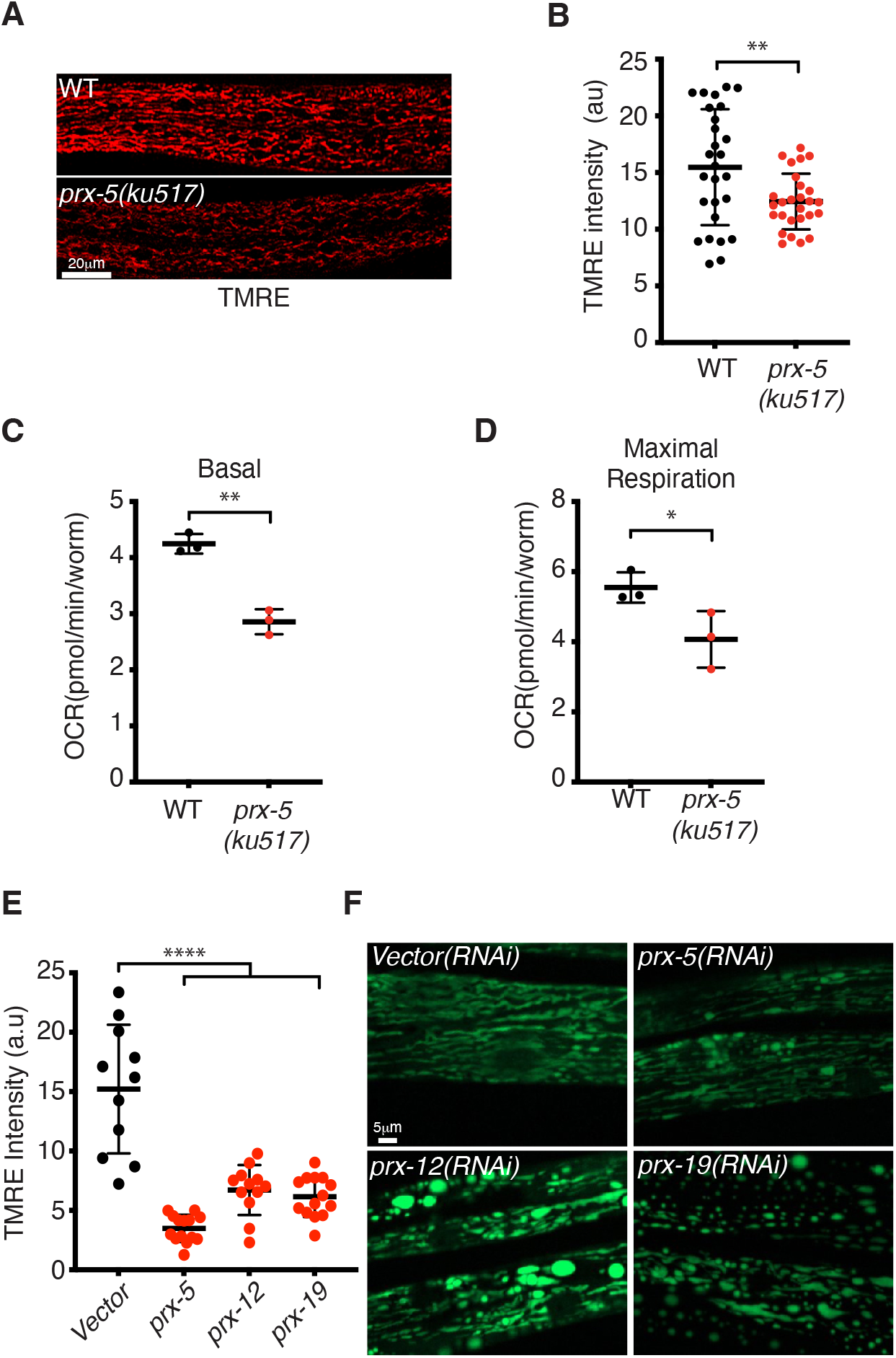
Peroxin perturbations lead to mitochondria dysfunction. **a.** TMRE staining of wildtype and *prx-5*(*ku517*) worms. Scale bar 20 μm. Experiments were repeated 3 times with similar results. **b.** Quantification of TMRE intensity in the intestine of wildtype and *prx-5*(*ku517*) worms. Experiments were repeated 3 biologically independent times with similar results. N=21 worms (wildtype), N=33 worms *prx-5*(*ku517*)). Error bars mean +/− s.d, (two tailed Student’s *t*-test). a.u. – Arbitrary units. **c-d.** Oxygen consumption rates (OCR) in wildtype and *prx-5*(*ku517*) worms. Basal respiration (**c**), maximal respiration (**d**). Error bars mean +/− s.d, (two tailed Student’s *t*-test). Experiment was performed 3 biologically independent times with similar results. **e.** Quantification of TMRE intensity in the intestine of wildtype worms treated with control, *prx-5, prx-12* or *prx-19* RNAi. Experiments were repeated 3 biologically independent times with similar results. N=11 (control),14 (*prx-5*), 12 (*prx-12*), 14 (*prx-19*). Error bars mean +/− s.d, (two tailed Student’s *t*-test). a.u. – Arbitrary units. **f.** Worms expressing *myo-3_pr_-:: ^mt^gfp* treated with control, *prx-5, prx-12* or *prx-19* RNAi. Scale bar 5 μm.

Since mitochondria dysfunction typically lead to the induction of the UPR^mt^^27^, we hypothesized that defects in peroxins will result in UPR^mt^ activation. Using the UPR^mt^ reporter strain *hsp-6_pr_::gfp*, which measure the transcription of the conserved mitochondrial chaperone *hsp-6*^28^, we compared the intensity of GFP in WT worms and in the *prx-5*(*ku517*) mutant strain. As depicted in Fig. 2*a*, *hsp-6* was induced in the *prx-5* mutant in an *atfs-1* dependent manner. Moreover, gene expression analysis indicated that *atfs-1* was induced in the *prx-5* loss of function strain as well as *hsp-6* (albeit insignificantly) (Fig 2*b*). Induction of the UPR^mt^ was not limited to *prx-5* and was observed when other peroxins were targeted via RNAi (Fig 2*c*). Together, these results indicate that defects in peroxisome biogenesis cause mitochondria stress which leads to the induction of the UPR^mt^.

**Figure 2:**
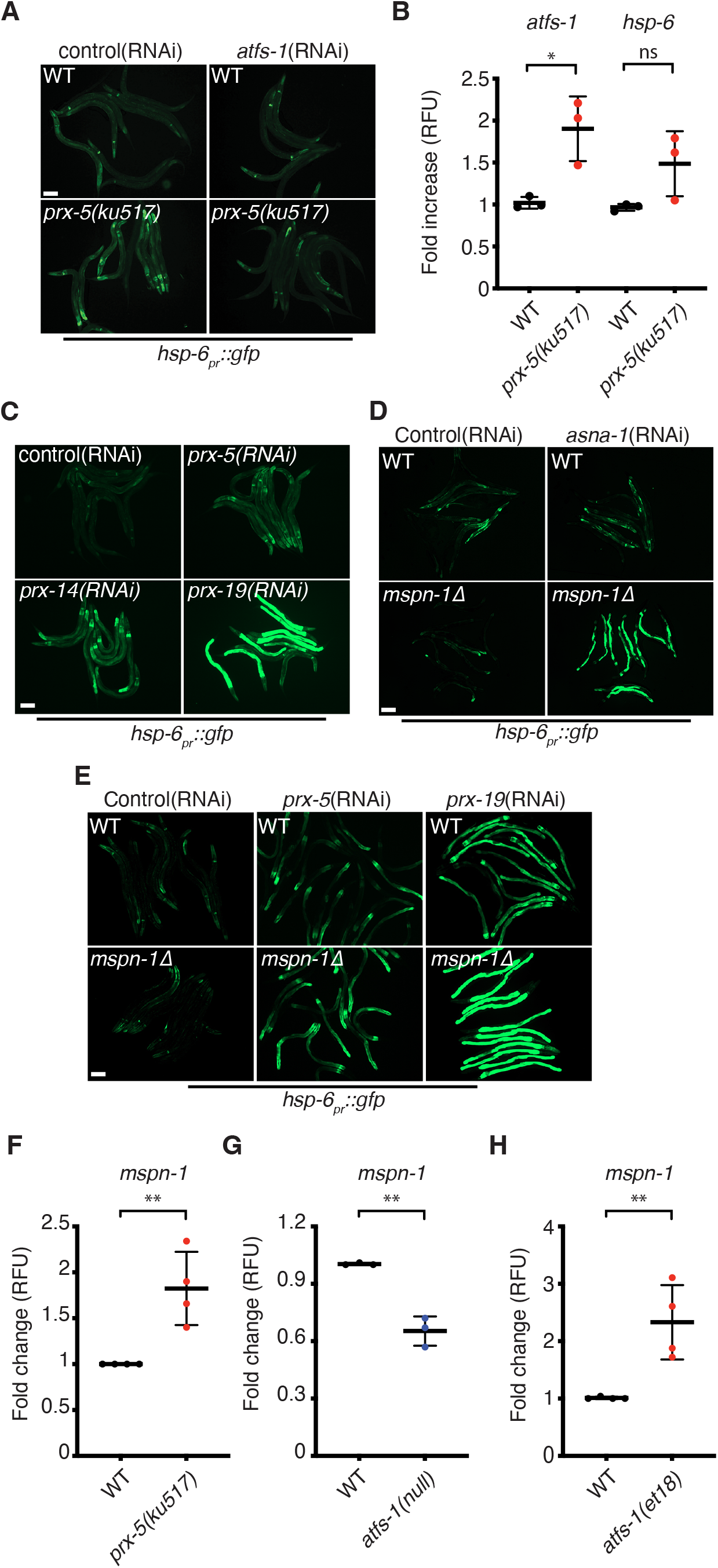
Peroxin defects lead to protein mislocalization and induce the UPR^mt^. **a.** *hsp-6_pr_::gfp* in wildtype and *prx-5*(*ku517*) worms treated with control or *atfs-1* RNAi. Scale bar 0.1 mm. N=3 biologically independent experiments with similar results. **b.** Transcript levels of activated transcription factor stress-1 (*atfs-1*) and of heat shock protein-6 (*hsp-6*) as determined by qRT-PCR in wildtype and *prx-5*(*ku517*) worms. N=3 biologically independent experiments. Error bars mean +/− s.d, (two tailed Student’s *t*-test). RFU – relative fluorescence units. **c.** *hsp-6_pr_::gfp* in worms raised on control, *prx-5, prx-14 and prx-19* RNAi. Scale bar 0.1 mm. Experiments were repeated 3 biologically independent times with similar results. **d.** *hsp-6_pr_::gfp* in wildtype and Δ*mspn-1* worms treated with control or *asna-1* RNAi. Scale bar 0.1 mm. N=3 biologically independent experiments with similar results. **e.** *hsp-6_pr_::gfp* in wildtype and Δ*mspn-1* worms treated with control, *prx-5* or *prx-19* RNAi. Scale bar 0.1 mm. N=3 biologically independent experiments with similar results. **f.** Transcript levels of *mspn-1* as determined by qRT-PCR in wildtype and *prx-5*(*ku517*)worms. N=4 biologically independent experiments. Error bars mean +/− s.d, (two tailed Student’s *t*-test). RFU – relative fluorescence units. **g.** Transcript levels of *mspn-1* as determined by qRT-PCR in wildtype and *atfs-1*(*null*)worms. N=3 biologically independent experiments. Error bars mean +/− s.d, (two tailed Student’s *t*-test). RFU – relative fluorescence units. **h.** Transcript levels of *mspn-1* as determined by qRT-PCR in wildtype and *atfs-1(et18*) worms. N=4 biologically independent experiments. Error bars mean +/− s.d, (two tailed Student’s *t*-test). RFU – relative fluorescence units.

### The UPR^mt^ regulates the transcription of *mspn-1*

We next considered the cause for mitochondrial dysfunction in the scenario of peroxin defects. It was recently shown in yeasts and mammals that peroxisomal proteins mislocalize to the mitochondria during peroxin deficiencies and cause mitochondrial dysfunction^11^. To alleviate the stress, cells utilize the mitochondrial outer membrane translocase ATAD1/MSPN-1, which removes mislocalized proteins from the mitochondria^11^. MSPN-1 shares strong sequence similarities to the mammalian ATAD1 and is the worm homolog of ATAD1 (Fig. S1*a*). To investigate the cause for mitochondrial dysfunction we created a deletion strain of *mspn-1* using CRISPR-Cas9. This deletion lacks the entire functional AAA domain of *mspn-1*^29^ and the strain was named Δ*mspn-1* (Fig. S1*b*). To determine whether the function of ATAD1/MSPN-1 is conserved in *C.elegans*, we mimicked conditions that promote protein mislocalization. Deletion of components required for the ER tail-anchored targeting pathway (GET complex), results in mislocalization of proteins to mitochondria^30^ and the consequential extraction by ATAD1/MSPN-1 in yeasts and mammals^12,13^. We reasoned that the inability to extract mislocalized proteins by MSPN-1 will result in mitochondrial stress that will activate the UPR^mt^.

Deletion of *mspn-1* alone did not lead to the activation of the UPR^mt^, suggesting that under basal conditions protein mislocalization is mild and has little effect on the integrity of the mitochondria. Similarly, knockdown of the GET complex subunit get3/*asna-1* in WT worms had little effect on the activation of the UPR^mt^. However, when combined with the Δ*mspn-1* strain, the UPR^mt^ was strongly induced (Fig. 2*d*). These results indicate that *mspn-1* and *asna-1* have negative genetic interactions, as was reported in yeasts and mammals and supports the role of MSPN-1 in extracting mislocliazed proteins from the mitochondria^11–13^. Next, we knockdown *prx-19* and *prx-5* in WT and Δ*mspn-1* worms. Knockdown of the peroxins led to induction of the UPR^mt^, which was exacerbated in the Δ*mspn-1* strain (Fig. 2*e*). These results indicate that MSPN-1 alleviates mitochondria stress during peroxin deficiencies.

Importantly, the expression of *mspn-1* was upregulated in the *prx-5*(*ku517*)mutant strain (Fig. 2*f*), suggesting that cells elicit a protective response against peroxisomal stress. Since the UPR^mt^ is induced during peroxisomal perturbations, we tested whether it regulates the expression of *mspn-1*. To this end we utilized strains that constitutively activate (*et18*) or lack ATFS-1, the transcription factor that is responsible for the induction of the UPR^mt^. *mspn-1* transcripts were upregulated in the *atfs-1*(*et18*) strain while downregulated in the *atfs-1*(*null*) strain, indicating that *mspn-1* expression is regulated by the UPR^mt^ (Fig. 2*g*-*h*). Together, our results suggest that protein mislocalization contributes to mitochondrial stress during peroxin deficiencies and to the induction of the UPR^mt^, which in turn regulates the expression of *mspn-1* to alleviate the stress.

### The UPR^mt^ regulates genes required for peroxisome assembly

The UPR^mt^ is a broad transcriptional program that extends beyond the production of proteases and chaperones^14,17^. Upon analysis of previous deep sequencing results of constitutively active and null *atfs-1* strains we found that out of the 12 peroxin genes in *C. elegans*, 6 were significantly upregulated when ATFS-1 was constitutively active and 9 were downregulated in the *atfs-1*(*null*) strain (Fig. 3*a* and S2*a*)^17^. We validated the expression of several conserved subunits that are essential for peroxisome biogenesis, including *prx-5*, which encodes a protein essential for importing proteins to the peroxisome matrix^23,24^ as well as *prx-3* and *prx-19* that are required for the transport of peroxisomal membrane proteins^31,32^. We also tested the expression of the peroxisomal fatty acid transporter *pmp-4*, an orthologue of human ABCD1 whose mutations lead to the most frequent peroxisomal disorder, X-linked adrenoleukodystrophy ^33^ and that was found in our deep sequencing to be downregulated in the *atfs-1*(*null*) strain (Fig. S2*a*). As depicted in Figs. 3*b*–*3c* all genes were upregulated in the *atfs-1(et18*) strain and excluding *prx-19* all were significantly downregulated in the *atfs-1*(*null*) strain. Importantly, a previous ChIP-seq analysis of ATFS-1, identified an interaction with the promoter of *prx-19*^18^, suggesting that ATFS-1 directly regulates peroxisome gene expression. To validate this observation, we performed ChIP using ATFS-1 antibodies followed by qPCR for the promoter of *prx-19*. Indeed, ATFS-1 bound efficiently to the *prx-19* promoter (Fig. 3*d*). Together, our results indicate that ATFS-1 regulates genes required for peroxisome assembly.

**Figure 3:**
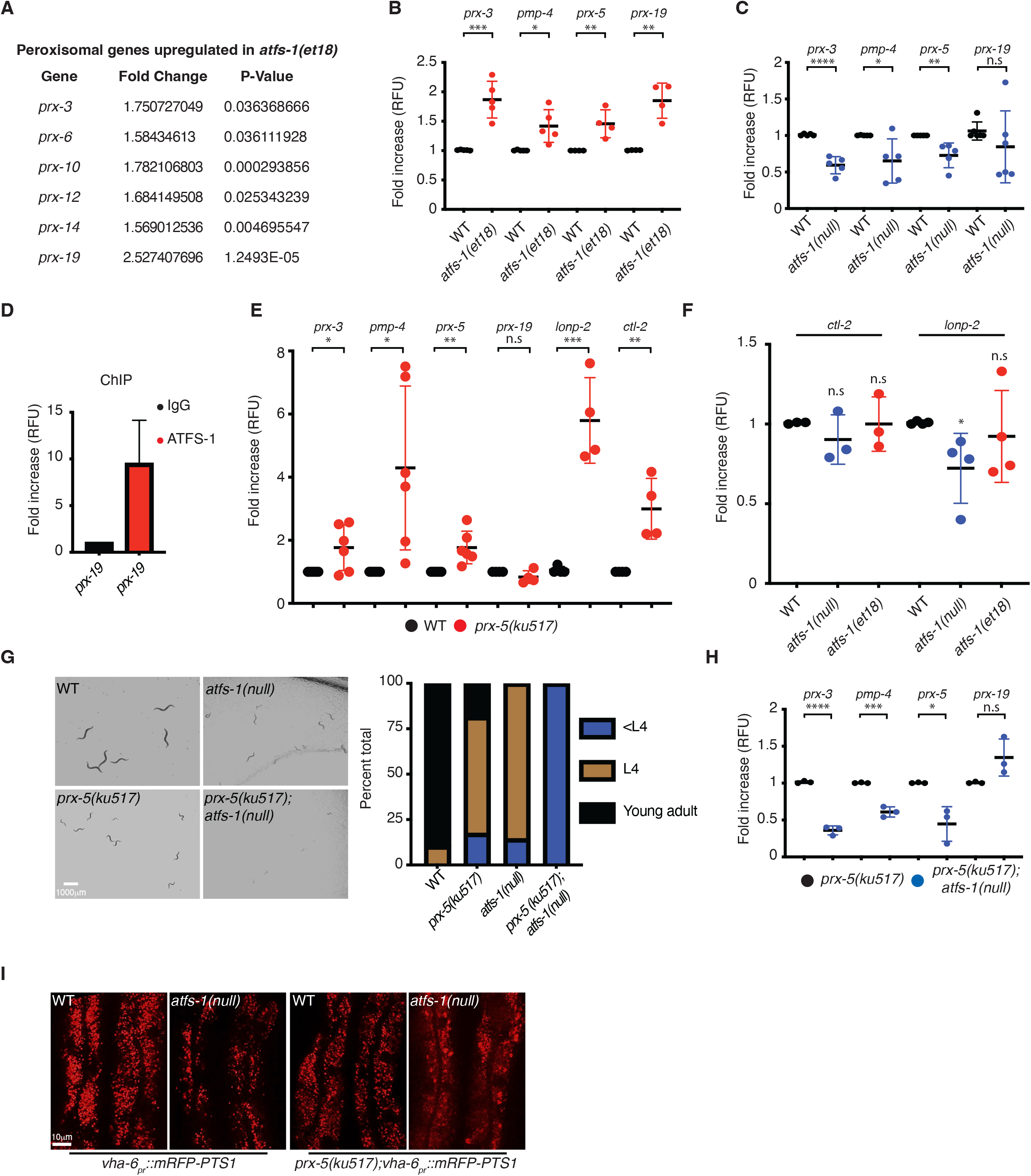
ATFS-1 regulates peroxisome biogenesis. **a.** Deep sequencing results depicting peroxin genes regulated by constitutive activation of ATFS-1. **b.** Transcript levels of *prx-3, pmp-4, prx-5 and prx-19* as determined by qRT-PCR in wildtype and *atfs-1(et18*) worms. N=5 (*prx-3, pmp-4*), N=4 (*prx-5, prx-19*) biologically independent experiments. Error bars mean +/− s.d, (two tailed Student’s *t*-test). RFU – relative fluorescence units. **c.** Transcript levels of *prx-3, pmp-4, prx-5 and prx-19* as determined by qRT-PCR in wildtype and *atfs-1*(*null*) worms. N=5 biologically independent experiments. Error bars mean +/− s.d, (two tailed Student’s *t*-test). RFU – relative fluorescence units. **d.** ChIP of *prx-19* promoter in wildtype worms as measured by qRT-PCR. N=2. biologically independent experiments. RFU – relative fluorescence units. **e.** Transcript levels of *prx-3, pmp-4, prx-5, prx-19, ctl-2 and lonp-2* as determined by qRT-PCR in wildtype and *prx-5*(*ku517*) worms. N=6 (*prx-3, pmp-4, prx-5*) n=4 (*prx-19, ctl-2, lonp-2*) biologically independent experiments. Error bars mean +/− s.d, (two tailed Student’s *t*-test). RFU – relative fluorescence units. **f.** Transcript levels of *ctl-2 and lonp-2* as determined by qRT-PCR in wildtype, *atfs-1*(*null*)and *atfs-1(et18*) worms. N=3. (*ctl-2*) N=4 (*lonp-2*) biologically independent experiments. Error bars mean +/− s.d, (two tailed Student’s *t*-test). RFU – relative fluorescence units. **g.** Representing photomicrographs of wildtype, *atfs-1(null), prx-5*(*ku517*) and *prx-5(ku517);atfs-1*(*null*) worms raised on *E.coli* for 3 days. Scale bar 1mm (left). And quantification of worm development (right). N=279 (wildtype), *249(prx-5(ku517)*), 210 (*atfs-1(null*)) and *152(prx-5(ku517);atfs-1*(*null*)). **h.** Transcript levels of *prx-3, pmp-4, prx-5 and prx-19* as determined by qRT-PCR in *prx-5*(*ku517*) and *prx-5(ku517);atfs-1*(*null*) worms. N=3 biologically independent experiments. Error bars mean +/− s.d, (two tailed Student’s *t*-test). RFU – relative fluorescence units. **i.** *vha-6pr::mRFP-pts1* in wildtype, *atfs-1(null), prx-5*(*ku517*) and *prx-5(ku517);atfs-1*(*null*) worms. Scale bar 0.1 mm. Experiments were repeated 3 biologically independent times with similar results.

### ATFS-1 regulates the peroxisomal retrograde response and is essential for worm development during peroxisomal stress

Next, we considered whether ATFS-1 participates in regulating peroxisomal transcripts during peroxin perturbations. Knockdown of *prx-5* by RNAi was recently shown to induce a retrograde response, which leads to the induction of numerous peroxisomal genes, including peroxins, peroxisomal transporters, the peroxisomal catalase (*ctl-2*)and the peroxisomal Lon protease (*lonp-2*)^34^. In support of these findings, we found that *lonp2, ctl-2*, peroxins and *pmp-4* were upregulated in the *prx-5*(*ku517*) strain (Fig. 3*e*). However, while constitutive activation of ATFS-1 led to the induction of *prx-19* (Fig. *3b*),it remained unchanged in the *prx-5*(*ku517*) mutant strain. Moreover, *atfs-1(et18*) did not significantly alter the transcript levels of *ctl-2* and *lonp-2*, though *lonp-2* was downregulated in the *atfs-1*(*null*) strain (Fig 3*f*). Therefore, activation of ATFS-1 alone doesn’t account for the entire peroxisomal retrograde response. To determine the role of ATFS-1 during peroxin perturbations we created a *prx-5(ku517); atfs-1*(*null*) double mutant strain. Strikingly, the *prx-5(ku517); atfs-1*(*null*) double mutant exhibited a strong negative interaction as the worms had severe developmental delays (Fig 3*g*) and led to reductions in the transcript levels of *prx-3, prx-5* and *pmp-4* but not of *prx-19* (Fig 3*h*).

Next, we monitored peroxisome morphology using a reporter strain in which the peroxisomal targeting sequence (pts1) was fused to mRFP^35^. mRPF-pts1 exhibited a punctuated phenotype typical to peroxisomes in wildtype, *atfs-1*(*null*) and in the *prx-5*(*ku517*) mutant strains (Fig 3*i*). However, combining *atfs-1*(*null*) with the *prx-5*(*ku517*) mutant strain resulted in the cytosolic accumulation of mRFP and in the appearance of large peroxisomes (Fig 3*i*). Together, our results indicate that ATFS-1 regulates peroxisomal genes and is required for worm development and peroxisome maintenance during peroxisomal stress.

### NHR-49 genetically interact with ATFS-1 and is a key component of the peroxisomal retrograde response

Since constitutive activation of ATFS-1 did not lead to the induction of *ctl-2* and *lonp-2*, which were otherwise upregulated in the *prx-5*(*ku517*) mutant strain, we sought to identify additional transcription factors that are involved in peroxisome biogenesis. NHR-49 is a functional homolog of the human peroxisome proliferator activator receptor alpha (PPARα)^22^ and was recently shown to regulate the transcripts of *ctl-2* and *lonp-2* during *prx-5* knockdown^34^. We thus tested the role of *nhr-49* in the *prx-5* mutant strain.

Knockdown of *nhr-49* in the *prx-5*(*ku517*) mutant strain resulted in marked transcriptional reductions of *ctl-2, lonp-2*, the peroxisomal transporter *pmp-4* as well as the peroxins *prx-3* and *prx-5*, while *prx-19* remained unaffected (Fig 4*a*). Thus, *nhr-49* knockdown resulted in the downregulation of all the tested genes that were induced in the *prx-5*(*ku517*) mutant strain (Fig 3*e*). Importantly, knockdown of *nhr-49* in the background of the *prx-5*(*ku517*) mutant resulted in the upregulation of *atfs-1* transcripts as well as of *hsp-6* (albeit insignificantly) (Fig 4*b*), pointing to a feedback loop between NHR-49 and ATFS-1 and suggesting that mitochondrial stress increased in the absence of *nhr-49*.

**Figure 4:**
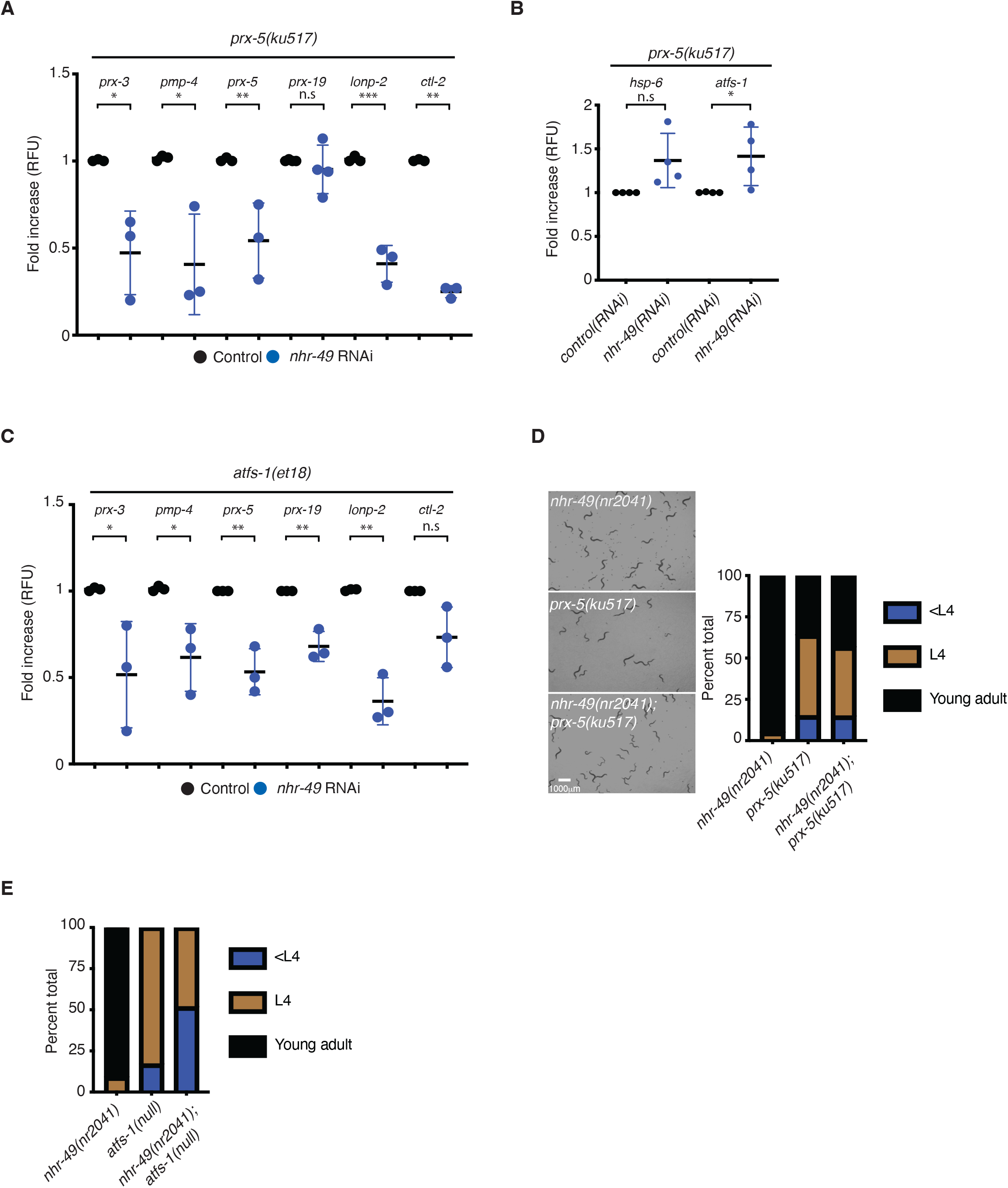
*nhr-49* regulates the peroxisomal retrograde response and genetically interacts with *atfs-1*. **a.** Transcript levels of *prx-3, pmp-4, prx-5, prx-19, lonp-2 and ctl-2* as determined by qRT-PCR in *prx-5*(*ku517*) worms treated with control or *nhr-49* RNAi. N=3 biologically independent experiments. Error bars mean +/− s.d, (two tailed Student’s *t*-test). RFU – relative fluorescence units. **b.** Transcript levels of *hsp-6* and *atfs-1* as determined by qRT-PCR in *prx-5*(*ku517*) worms treated with control or *nhr-49* RNAi. N=4 biologically independent experiments. Error bars mean +/− s.d, (two tailed Student’s *t*-test). RFU – relative fluorescence units. **c.** Transcript levels of *prx-3, pmp-4, prx-5, prx-19, lonp-2 and ctl-2* as determined by qRT-PCR in *atfs-1(et18*) worms treated with control or *nhr-49* RNAi. N=3 biologically independent experiments. Error bars mean +/− s.d, (two tailed Student’s *t*-test). RFU – relative fluorescence units. **d.** Representing photomicrographs of *nhr-49(nr2041), prx-5*(*ku517*) and *nhr-49(nr2041);prx-5*(*ku517*) worms raised on *E.coli* for 3 days. Scale bar 1mm (left). And quantification of worm development (right). N=373 (*nhr-49(nr2041)*), 326 (*prx-5*(*ku517*))and 384 (*nhr-49(nr2041);prx-5(ku517)*). **e.** Worm development Quantification of *nhr-49(nr2041), atfs-1*(*null*) and *nhr-49(nr2041);atfs-1*(*null*) worms raised on *E.coli* for 3 days. N=224 (*nhr-49(nr2041)*), 202 (*atfs-1(null*)) and 190 (*nhr-49(nr2041);atfs-1*(*null*)).

Next, we assessed the role of *nhr-49* upon constitutive activation of ATFS-1. Knockdown of *nhr-49* in the *atfs-1(et18*) mutant strain resulted in a marked reduction of all the peroxisomal genes tested (Fig 4*c*). Taken together, our results indicate that *nhr-49* is a key component in regulating peroxisomal transcripts.

Since *nhr-49* play key roles in the induction of peroxisomal transcripts, we considered the genetic interactions between *nhr-49, atfs-1* and *prx-5*. To this end we utilized an *nhr-49* null strain (*nhr-49(nr2041*))^22^ and crossed it with *prx-5*(*ku517*) or with *atfs-1(null*). In contrast to the *prx-5(ku517);atfs-1*(*null*) double mutant that exhibited severe developmental delays (Fig 3*g*), the *prx-5(ku517);nhr-49(nr2041*) double mutant did not alter developmental (Fig 4*d*). This suggests that the role of ATFS-1 in protecting the mitochondria during peroxisomal stress is key to worm development and that ATFS-1 may compensate for the loss of *nhr-49*. In view of this, a combination of *nhr-49* and *atfs-1* null mutants resulted in developmental delays (Fig 4*e*). Together, our results indicate a negative genetic interaction between the two transcription factors, supporting the notion that ATFS-1 activity is essential during reduced peroxisomal function.

## Discussion

Peroxisomes dysfunction is associated with the aging process and numerous diseases including metabolic disorders and neurodegeneration. Despite this, little is known about protective responses that cells employ to protect peroxisomes. Here we show that ATFS-1 elicits an indispensable response that is essential for peroxisome assembly and worm development.

In mammals, the transcriptional coactivator PGC-1α acts as a master regulator for mitochondria and peroxisome biogenesis^36–38^. Worms lack PGC-1α, however ATFS-1 shares many of its functions and acts as a biogenesis regulator for both organelles. This study highlights a potential coordinated biogenesis response of peroxisomes and mitochondria; Peroxisome perturbations result in mitochondria stress that activates the UPR^mt^, which induces genes that are essential for the biogenesis of both organelles.

Importantly, ATFS-1 was previously shown to regulate the transcription of fission factors^14,17^. Fission factors are shared between mitochondria and peroxisomes and allow preexisting organelles to multiply^39^. We observed large peroxisomal structures during peroxisomal stress in the *atfs-1*(*null*) strain and the accumulation of cytosolic mRFP-pts1. This may reflect a combination of reduced fission activity and transcription of peroxins orchestrated by ATFS-1.

We have further identified that the UPR^mt^ regulates the transcription of the AAA-ATPase ATAD1/MSPN-1, which extracts mislocalized proteins from the mitochondria. Protein mislocalization plays a major role in mitochondrial damage during peroxin deletion in yeasts and mammals and overexpression of ATAD1 can alleviate the stress^11^. It was thus important to identify how the transcription of MSPN-1 is regulated. And while protein mislocalization was not shown in *C.elegans*, our genetic data is consistent with the results obtained from yeasts and mammals and points to a conserved mechanism^11–13^.

Constitutive activation of ATFS-1 does not account for the entire retrograde response elicited in the *prx-5* mutant. Since *nhr-49* was previously shown to regulate *ctl-2* and *lonp-2*^34^, we investigated its role during peroxisomal stress. Knockdown of *nhr-49* accounted for all the peroxisomal genes that were induced in the *prx-5* and *atfs-1(et18*) mutants, suggesting that it’s indispensable for the peroxisomal retrograde response. However, unlike the *atfs-1*(*null*) strain, deletion of *nhr-49* combined with *prx-5* mutant did not result in developmental delays. Since ATFS-1 promotes mitochondrial biogenesis, the developmental delays in the *atfs-1*(*null*) can be attributed to decreased mitochondrial function. However, since ATFS-1 regulates mitochondrial genes that are shared with peroxisomes, the impact on peroxisomes may be more severe in the absence of *atfs-1*. Lastly, we have identified genetic interactions and a feedback loop between ATFS-1 and NHR-49. And while it is unknown how NHR-49 is activated during peroxisomal stress, it is plausible that ATFS-1 regulates a subset of PPARs and NHR-49 to promote peroxisome biogenesis.

## Methods

### Worms, plasmids and staining

The reporter strain *hsp-6_pr_::gfp* for visualizing UPR^mt^, the mRFP-pts1 to visualize peroxisomes and the *myo-3_pr_::^mt^gfp* for visualization of mitochondrial mass have been previously described^28,35,40^. The *atfs-1(et18*) strain was a gift from Marc Pilon. N2(wildtype), *prx-5 (ku517*) and *nhr-49(nr2041*) were obtained from the Caenorhabditis Genetics Center (Minneapolis, MN). The *mspn-1* Δ strain was generated via CRISPR-Cas9 in N2 worms. The crRNAs (IDT) were co-injected with purified Cas9 protein, tracrRNA (Dharmacon), repair templates (IDT) and the pRF4::rol-6(su1006) plasmid as described^41,42^. The crRNAs and repair templates used in this study are listed in Table 1. The pRF4::rol-6 (su1006) plasmid was a gift from Craig Mello^43^. Worms were raised HT115 strain of *E. coli* and RNAi performed as described^44^. TMRE experiments were performed by synchronizing and raising worms on plates previously soaked with M9 buffer containing EtBr or 2μM TMRE. Worms were analyzed at the L4 larvae stage

**Table 1.**
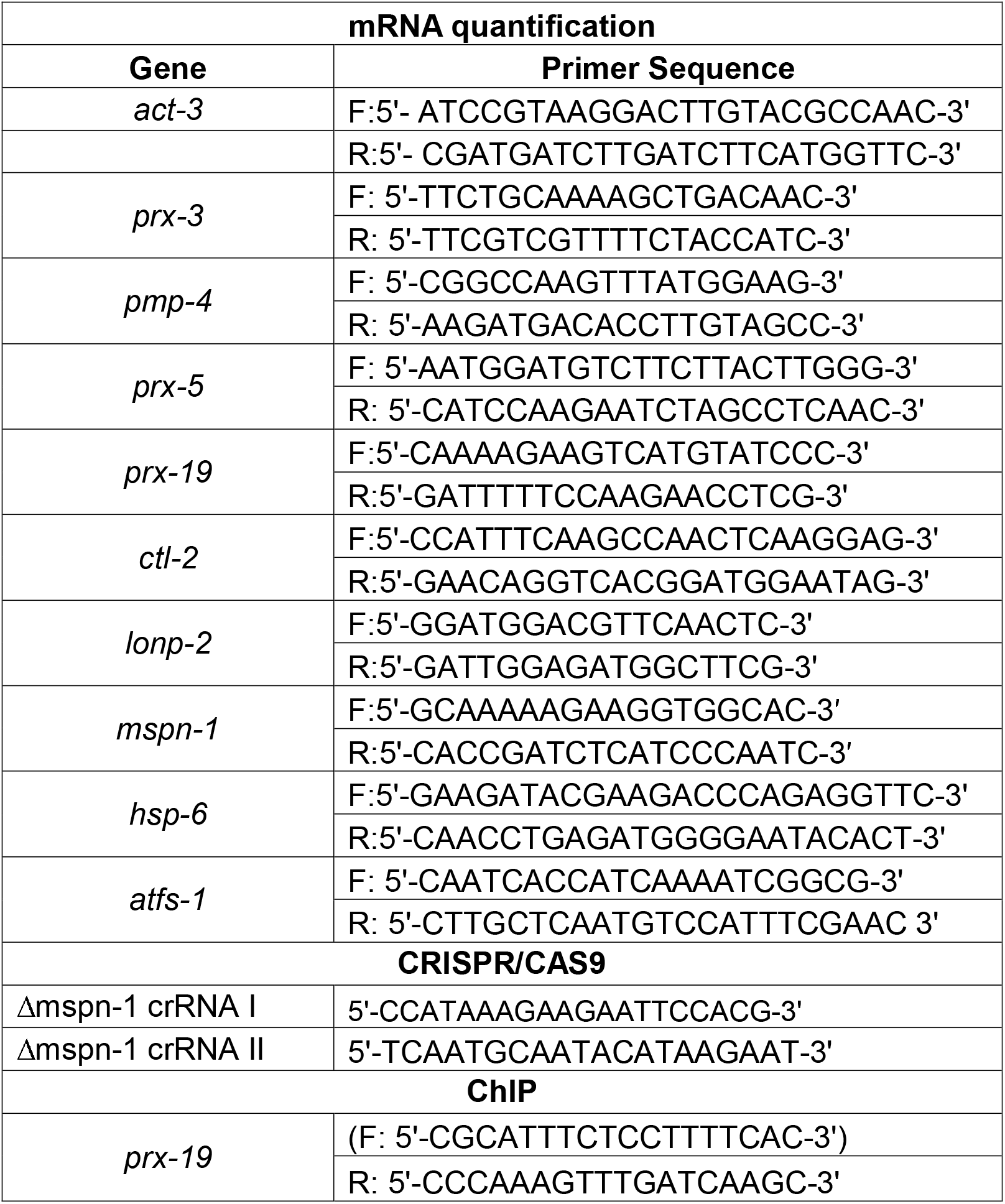
Primers used in this study

### RNA isolation and qRT-PCR

RNA isolation and qRT-PCR analysis were previously described^20^. Worms were synchronized by bleaching, raised on HT115 *E. coli* and harvested at the L4 stage. Total RNA was extracted from frozen worm pellets using RNA STAT (Tel-Test) and 500 ng RNA was used for cDNA synthesis with qScript^™^ cDNA SuperMix (QuantaBio). qPCR was performed using iQ^™^ SYBR^®^ Green Supermix (Bio-Rad Laboratories). qPCR primers are listed in Table 1. All qPCR results were repeated at least 3 times and performed in triplicates. A two tailed Student’s *t*-test was employed to determine the level of statistical significance.

### Oxygen Consumption

Oxygen consumption was measured using a Seahorse XFe96 Analyzer at 25°C as described previously^45^. In brief, L4 worms were transferred onto empty plates and allowed to completely digest the remaining bacteria for 1 hour, after which 10 worms were transferred into each well of a 96-well microplate containing 180 μl M9 buffer. Basal respiration was measured for a total of 30 minutes, in 6 minute intervals that included a 2 minute mix, a 2 minute time delay and a 2 minute measurement. To measure respiratory capacity, 15 μM FCCP was injected, the OCR (oxygen consumption rate) reading was allowed to stabilize for 6 minutes then measured for five consecutive intervals. Mitochondrial respiration was blocked by adding 40mM Sodium azide. Each measurement was considered one technical replicate.

### Analysis of worm development

Worms were synchronized via bleaching and allowed to develop on HT115 bacteria plates for 3 days at 20°C. Developmental stage was quantified as a percentage of the total number of animals. Each experiment was preformed three times.

### Statistics

All experiments were performed at least three times yielding similar results and comprised of biological replicates except for ChIP that was preformed two times. The sample size and statistical tests were chosen based on previous studies with similar methodologies and the data met the assumptions for each statistical test performed. No statistical method was used in deciding sample sizes. Blinded analyses were performed for developmental and TMRE quantifications, and randomization was not used. For all figures, the mean ±standard deviation (s.d.) is represented unless otherwise noted. Prism 9 (GraphPad) is used for statistical analysis and graph creation.

### Microscopy

*C. elegans* were imaged using either a Zeiss AxioCam 506 mono camera mounted on a Zeiss Axio Imager Z2 microscope or a Zeiss AxioCam MRc camera mounted on a Zeiss SteREO Discovery.V12 stereoscope. Images with high magnification (63×) were obtained using the Zeiss ApoTome.2. Exposure times were the same in each experiment.

### Chromatin immunoprecipitation (ChIP)

ChIP assays for ATFS-1 were performed as previously described^18^. Synchronized worms were cultured in liquid and harvested at early L4 stage by sucrose flotation. The worms were lysed via Teflon homogenizer in cold PBS with protease inhibitors (Roche). Cross-linking of DNA and protein was performed by treating the worms with 1.85% formaldehyde with protease inhibitors for 15 min. Glycine was added to a final concentration of 125 mM and incubated for 5 min at room temperature to quench the formaldehyde. The pellets were resuspended twice in cold PBS with protease inhibitors. Samples were sonicated in a Bioruptor (Diagenode) for 15 min at 4°C on high intensity (30s on and 30s off). Samples were transferred to microfuge tubes and spun at 15,000*g for 15 min at 4°C. The supernatant was precleaned with pre-blocked ChIPgrade Pierce^™^ magnetic protein A/G beads (Thermo Scientific) and then incubated with ATFS-1 antibody or Mouse mAb IgG1 Isotype Control (Cell Signaling Technology, G3A1) rotating overnight at 4°C. The antibody-DNA complex was precipitated with protein A/G magnetic beads or protein A sepharose beads (Invitrogen). After washing, the crosslinks were reversed by incubation at 65°C overnight. The samples were then treated with RNaseA at 37°C for 1.5 hour followed by proteinase K at 55°C for 2 hours. Lastly, the immunoprecipitated and input DNA were purified with ChIP DNA Clean & Concentrator (Zymo Research, D5205) and used as templates for qPCR.

## Notes

### Competing Interest Statement

The authors have declared no competing interest.

