## Supplementary material for "ATFS-1 regulates peroxisome assembly genes and protects both mitochondria and peroxisomes during peroxin perturbations": Figure S1

**A**

```
ATAD1           MVHAEAFSRPLSRNEVVGLIFRLTIFGAVTYFTIKWMVDAIDPTRKQKVEAQKQAEKLMK
MSPN-1          ---------MTDRNELIGVAIRVVAAAAVSFLSVRYLVKYLDPNYSVNEESKKKVAQLFH
                           .***::*: :*:.  .**:::::.::*. :**. . : *::*:. :*::

ATAD1           QIGV-KNVKLSEYEMSIAAHLVDPLNMHVTWSDIAGLDDVITDLKDTVILPIKKKHLFEN
MSPN-1          ELGIDRQIELSEHEIRIATQFVGGEDVGADWDEIGGCEELVAELKDRIILPLRFASQ-SG
                ::*: .:::***:*: **:::*.  :: . *.:*.* ::::::*** :***:.     ..

ATAD1           SRLLQPPKGVLLYGPPGCGKTLIAKATAKEAGCRFINLQPSTLTDKWYGESQKLAAAVFS
MSPN-1          SHLLSPPRGILLYGPPGCGKTLLAKAVARAAGCRFINLQVSNLTDKWYGESQKLAAAVFS
                *.**.**.*:************:***.*. ********* *.******************

ATAD1           LAIKLQPSIIFIDEIDSFLRNRSSSDHEATAMMKAQFMSLWDGLDTDHSCQVIVMGATNR
MSPN-1          VAQKFQPTIIFIDEIDSFLRDRQSHDHESTAMMKAQFMTLWDGFSSSGD-QIIVMGATNR
                :* *:**:************:*.* ***:*********:****:.:. . *:********

ATAD1           PQDLDSAIMRRMPTRFHINQPALKQREAILKLILKNENVDRHVDLLEVAQETDGFSGSDL
MSPN-1          PRDVDAAILRRMTARFQVPVPNAKQRSQILNVILRNEKINNTVNLGEIAQAAEGLSGSDL
                *.*:*:**:***.:**::  *  ***. **::**.**:::. *:* *:** ::*:*****

ATAD1           KEMCRDAALLCVREYVNSTSEESHDEDEIRPVQQQDLHRAIEKMKKSKDAAFQNVLTHVC
MSPN-1          KEVCR----LALLARAKATVASGGSVNQLLPLEQSDFESAVHKYMRA-----AHLLVEET
                **:**    *.:   .::*  .. . ::: *::*.*:  *: *  .:      ::*.

ATAD1           LD
MSPN-1          LD
                **
```

**B**

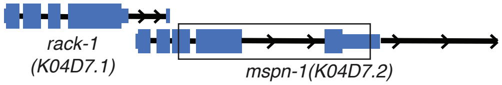
