## Supplementary material for "ATFS-1 regulates peroxisome assembly genes and protects both mitochondria and peroxisomes during peroxin perturbations": Figure S2

**A**

**Peroxisomal genes downregulated in *atfs-1(null)***

| Gene | Fold Change | P-Value |
|---|---|---|
| *prx-1* | 0.166346129 | 8.75662E-07 |
| *prx-2* | 0.330973411 | 0.040358656 |
| *prx-3* | 0.12478725 | 0.000425523 |
| *prx-5* | 0.251291637 | 0.001044767 |
| *prx-6* | 0.197785089 | 0.007226219 |
| *prx-11* | 0.213298624 | 2.87899E-05 |
| *prx-12* | 0.164314551 | 0.003552011 |
| *prx-13* | 0.218883786 | 0.00068612 |
| *prx-19* | 0.476700294 | 0.000943033 |
| *pmp-4* | 0.188269367 | 0.004556067 |
