## Supplemental Figure legend for "ATFS-1 regulates peroxisome assembly genes and protects both mitochondria and peroxisomes during peroxin perturbations"

**Figure S1: MSPN-1 sequence and deletion.**

**a.** Sequence alignment between the human ATAD1 and the worm MSPN-1. Sequence alignment was performed using Multiple Sequence Comparison by Log-Expectation (MUSCLE).

**b.** A schematic of *mspn-1(k04D7.2)* and *rack-1*(*k04D7.*1) genes. Box indicates the region that was deleted to create the Δ*mspn-1* strain.

**Figure S2: Peroxisomal genes downregulated in the *atfs-1(null)* strain.**

**a.** Deep sequencing results depicting genes regulated by deletion of ATFS-1.
